# Bantam regulates the adult sleep circuit in *Drosophila*

**DOI:** 10.1101/2021.08.21.457226

**Authors:** Michael Hobin, Katherine Dorfman, Mohamed Adel, Emmanuel J. Rivera-Rodriguez, Leslie C. Griffith

## Abstract

Sleep is a highly conserved feature of animal life characterized by dramatic changes in behavior, neural physiology and gene expression. The gene regulatory factors responsible for these sleep-dependent changes remain largely unknown. microRNAs are post-transcriptional modulators of gene expression which have been implicated in sleep regulation. Our previous screen identified 25 sleep-regulating microRNAs in *Drosophila melanogaster*, including the developmental regulator *bantam* (*ban*). Here we show that *ban* promotes early nighttime sleep through a population of glutamatergic neurons- the γ5β′2a/β′2mp/β′2mp_bilateral Mushroom Body Output Neurons (MBONs). We found that knockdown of *ban* in these neurons led to a reduction in early night sleep. The γ5β′2a/β′2mp/β′2mp_bilateral MBONs were previously shown to be wake-promoting, suggesting that *ban* acts to inhibit these neurons. GCaMP calcium imaging revealed that bantam inhibits the neural activity of the γ5β′2a/β′2mp/β′2mp_bilateral MBONs during the night but not the day. Blocking synaptic transmission in the γ5β′2a/β′2mp/β′2mp_bilateral MBONs rescued the effect of *ban* knockdown on sleep. Together these results suggest that *ban* promotes night sleep via the inhibition of the γ5β′2a/β′2mp/β′2mp_bilateral MBONs. RNAseq further revealed that bantam negatively regulates the wake-promoting mRNAs *Kelch* and *CCHamide-2 receptor* in the γ5β′2a/β′2mp/β′2mp_bilateral MBONs. These experiments establish *bantam* as an active regulator of sleep and neural activity within the fly brain.

## Introduction

Sleep is a behavioral state involving coordinated changes in gene expression and physiological activity across the brain and body. The decision to sleep is one that requires integration of information about both the internal state of the animal and its immediate external environment. Sleep circuitry has therefore evolved to have both dedicated and modulatory neuronal elements (Beckwith and French, 2019; Saper and Fuller, 2017; Shafer and Keene, 2021). Using multiple approaches, a still-expanding set of genes has also been linked to sleep (reviewed in Webb and Fu, 2020). A key challenge to the field of sleep is localizing the effect of a given sleep-regulating gene to a specific cell-type and physiological process.

Many of the identified sleep genes are involved in gene expression, including post-transcriptional regulators such as microRNAs. microRNAs are non-coding RNA transcripts approximately 22 bp in size that act to regulate mRNA expression (Huntzinger and Izaurralde, 2011). As part of the RNA-Induced-Silencing-Complex (RISC), microRNAs bind to complementary sequences known as microRNA Response Elements (MREs) on target mRNAs, generally within their 3’ UTRs. microRNA-mRNA binding reduces mRNA expression by promoting mRNA degradation or translational silencing. In the mammalian brain, microRNA expression is altered by sleep loss and expression of specific microRNAs induces changes in sleep and EEG (Davis et al., 2007; Davis et al., 2012; Davis et al., 2011) and multiple human sleep pathologies are associated with changes in microRNA expression (Holm et al., 2014).

In *Drosophila*, several microRNAs have been identified as regulators of sleep and circadian rhythmicity (Chen and Rosbash, 2017; Xia et al., 2020; Zhang et al., 2021) and this organism provides a useful system for modeling microRNA regulation of sleep. The fruit fly exhibits a homeostatically-regulated and circadian-timed diurnal sleep state characterized by a distinct posture, an increased arousal threshold and alterations in neural activity (Hendricks et al., 2000; Nitz et al., 2002; Shaw et al., 2000). The relatively simplified genome and nervous system of the fly as well as a versatile set of tools for their manipulation has allowed for the characterization of many sleep regulating genes and neural circuits (Shafer and Keene, 2021). To address the role of microRNAs in this behavior, we have used microRNA sponges, artificial RNA transcripts with multiple MREs for specific microRNAs to knockdown function (Fulga et al., 2015; Loya et al., 2009). Our recent screen of 143 microRNA sponges in *Drosophila* identified 25 sleep-regulating microRNAs-- 8 that promoted wake and 17 that promoted sleep (Goodwin et al., 2018). The latter group included *bantam* (*ban*), a microRNA which was initially identified as a growth promoting gene and has since been shown to be implicated in neurite growth and circadian rhythmicity (Jiang et al., 2014; Kadener et al., 2009; Lerner et al., 2015; Parrish et al., 2009). Although *ban* exhibits expression in multiple tissues throughout the lifespan of the fly, the cellular locus, target mRNA(s) and physiological basis of its sleep-regulating effects were unknown.

Here we map the nighttime sleep-promoting effect of *ban* to a subset of wake-promoting glutamatergic neurons- the γ5β′2a/β′2mp/β′2mp_bilateral MBONs (mushroom body output neurons). The mushroom body has been previously implicated in sleep, with each neuronal subtype in this structure having different functions (Aso et al., 2014b; Joiner et al., 2006; Pitman et al., 2006; Sitaraman et al., 2015b). *ban* promotes nighttime sleep by inhibiting the neural activity of the γ5β′2a/β′2mp/β′2mp_bilateral MBONs by downregulation of its wake-promoting target mRNAs *Kelch* (*kel*) and *CCHamide-2 receptor* (*CCHa2-R*).

## Results

In order to investigate the role of *ban* in sleep, we utilized a microRNA sponge transgene to reduce its levels (Fulga et al., 2015). We had previously shown that pan-neuronal (with *nSyb-GAL4*) expression of the *bantam* sponge *(ban-SP)* led to a reduction in sleep (Goodwin et al., 2018), indicating that *ban* acts in the nervous system to promote sleep. Since sleep-regulating genes can exert their effects at different stages throughout the lifespan of an animal, as developmental regulators contributing to the formation of sleep circuits (Chakravarti Dilley et al., 2020; Gong et al., 1998; Xie et al., 2019) or as active regulators of adult sleep (for review see Crocker and Sehgal, 2010) it was important to determine the temporal window of *ban* action.

We made use of an inducible expression system to control sponge expression. *ban-SP* was placed under the control of the pan-neuronal driver *nSyb-GAL4* and a ubiquitously expressed temperature-sensitive inhibitor of GAL4 activity-*tubulin-GAL80*^*ts*^ (McGuire et al., 2003). In this genotype, *ban-SP* is expressed at 29°C, but not at 17-18°C. Female *nSyb>ban-SP, tubulin-GAL80*^*ts*^ flies were raised at 17°C and then shifted to 29°C post-eclosion and for the duration of the sleep assay. Adult-specific pan-neuronal knockdown of *ban* led to a significant reduction in daytime and nighttime sleep compared to control animals expressing a scrambled sponge transgene (*UAS-scr-SP*) (Figure 1A). This reduction was most prominent in the early night, ZT12-ZT18 (Figure 1A). *nSyb>ban-SP, tubulin-GAL80*^*ts*^ flies also exhibited increased sleep-latency (time before sleep onset following lights off) (Supplementary Figure 1A). In contrast, developmental knockdown of *ban* led to a significant reduction in daytime, but not nighttime, sleep (Supplementary Figure 1B). These results indicate that in the adult brain, *ban* acts acutely to promote early night sleep.

**Figure 1.**
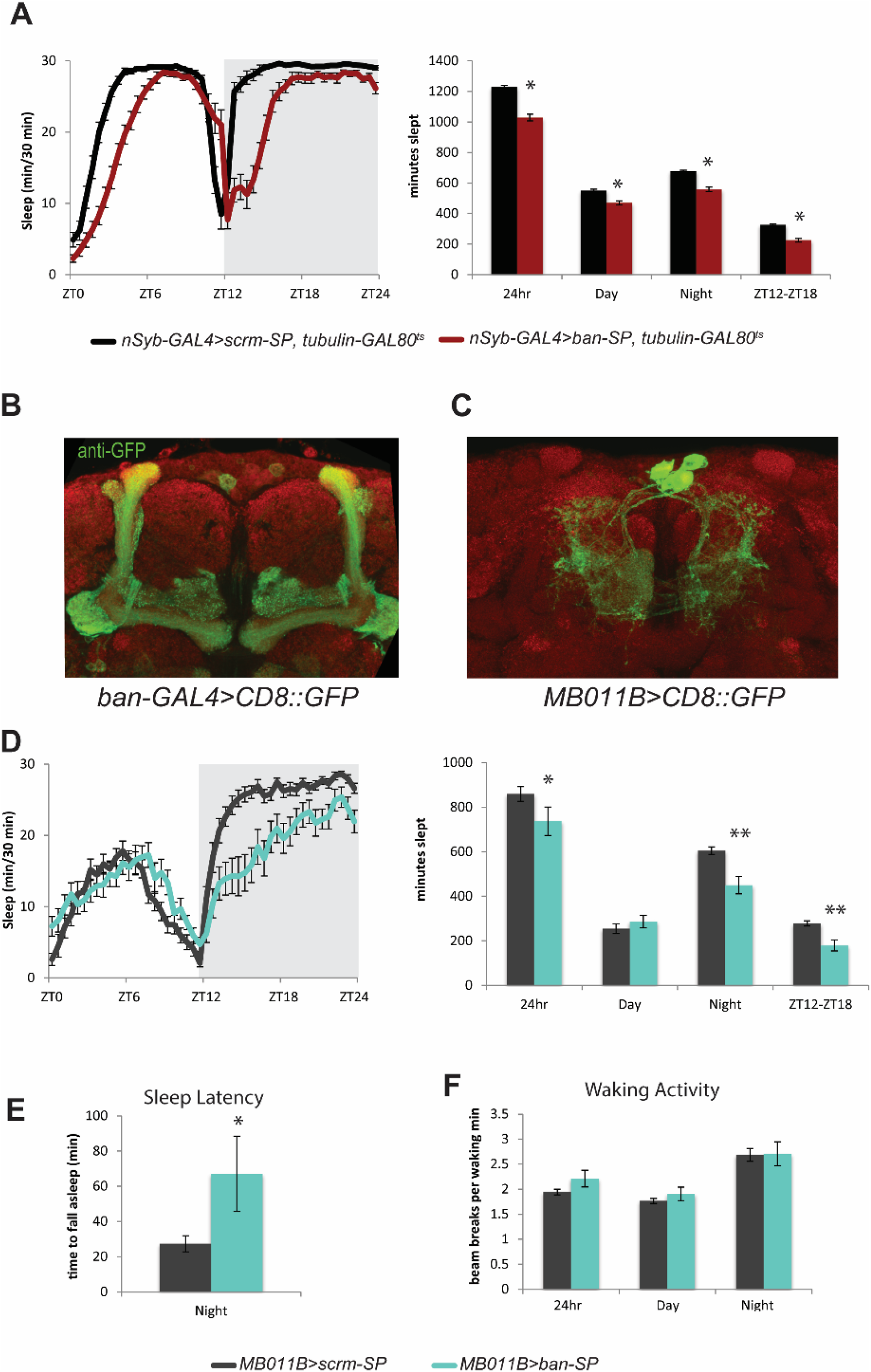
bantam promotes nighttime sleep via the γ5β′2a/β′2mp/β′2mp_bilateral MBONs (A) Adult knockdown of *ban*. Sleep data for *nSyb-GAL4>UAS-ban-SP, tubulin-GAL80*^*ts*^ and scramble controls raised at 17°C and tested at 29°C (mean± SEM). * represents P≤.0001 using the Kruskal-Wallis Test with Dunn’s multiple comparison test. (B) anti-GFP staining for the central brain of a representative *ban-GAL4>UAS-CD8::GFP* fly (63X). anti-BRP (nc82) used to stain neuronal anatomy (red). (C) anti-GFP and anti-BRP staining for the brain of a representative *MB011B>UAS-CD8::GFP* fly (63X). (D) Sleep data for *MB011B>UAS-ban-SP* and scramble controls (mean± SEM). * represents P≤.05 and ** represents P≤.0005 using the Mann-Whitney-Wilcoxon Test. (E) Sleep latency (minutes to fall asleep after night onset) for *MB011B>UAS-ban-SP* and scramble controls (mean± SEM). * represents P≤.05 for an unpaired t test. (F) Beam breaks per waking minute (general waking activity) for *MB011B>UAS-ban-SP* and scramble controls (mean± SEM).

We next sought to localize the effect of *ban* on sleep to a specific population of neurons. We expressed *CD8::GFP* under control of the *ban-GAL4* driver (generous gift of Sebastian Kadener) to examine the expression pattern of *ban* in the adult brain (Figure 1B). Immunohistochemistry of *ban>CD8::GFP* brains revealed high GFP staining in both the intrinsic and extrinsic cells of the mushroom body (MB; Figure 1B), a centrally located neuropil with a well-characterized role in *Drosophila* sleep (Joiner et al., 2006; Pitman et al., 2006). The mushroom body scaffold is composed of the axonal projections of Kenyon cells (intrinsic cells), which communicate with several populations of extrinsic cells. The primarily input cell type is dopaminergic (Mao and Davis, 2009), while the primary output of Kenyon cells is via synapses onto the dendrites of Mushroom Body Output Neurons (MBONs) (Aso et al., 2014a). Given the high expression level of *ban-GAL4* in the MB a/b core, we first examined whether *ban* expression in Kenyon cells was necessary for normal sleep. Expression of *ban-SP* under the powerful Kenyon cell driver *OK107-GAL4* had no effect on sleep (Supplementary Figure 2A), indicating that *ban* does not regulate sleep through MB intrinsic cells. We next tested the effect of *ban* knockdown on dopaminergic neurons. Expression of *ban-SP* under the dopaminergic driver *TH-GAL4* also had no effect on nighttime sleep, although it did lead to a small increase in daytime sleep (Supplementary Figure 2B). These results suggested that the cells in the *ban>CD8::GFP* staining pattern responsible for the early night sleep phenotype might be MBONs.

The MBONs consist of 21 subtypes divided into three classes determined by neurotransmitter identity (GABA, acetylcholine and glutamate) with each MBON named on the basis of the MB lobe section(s) which it innervates (Aso et al., 2014a). Visual inspection of *ban>CD8::GFP* brains revealed GFP-labeled MB extrinsic cells with centrally placed dorsal somata and strong innervation of the medial horizontal lobes (Figure 1B). The glutamatergic γ5β′2a/β′2mp/β′2mp_bilateral MBONs, which have a previously described role in sleep (Aso et al., 2014a; Sitaraman et al., 2015a), have a very similar morphology as shown by *MB011B>CD8::GFP* staining (Figure 1C), suggesting that *ban* is expressed in these neurons. In order to test if *ban* regulates sleep via this cell-type, we drove *ban-SP* with the *MB011B* split-GAL4. *MB011B>ban-SP* flies exhibited a significant reduction in total and nighttime sleep (Figure 1D), in addition to increased sleep latency (Figure 1E) compared to animals expressing a control scrambled sponge *(scr-SP*). Nighttime sleep loss was most severe in the early night ZT12-ZT18 period (Figure 1D). Interestingly, in contrast to the pan-neuronal adult knockdown results, *ban* knockdown in these MBONs had no effect on daytime sleep, indicating that the daytime sleep regulatory role of *ban* is likely mediated by different neurons. Importantly, knockdown of *ban* in the γ5β′2a/β′2mp/β′2mp_bilateral MBONs had no effect on waking motor activity (Figure 1F), demonstrating that the effect of this microRNA was specific to sleep rather than affecting general motor behavior.

microRNAs exert their effects on cellular functions via inhibition of target mRNAs either through transcript degradation or translational silencing (Jonas and Izaurralde, 2015). microRNA knockdown leads to increased expression of its target mRNAs, either through increased RNA stability, translation or both. Knockdown of *ban* in the γ5β′2a/β′2mp/β′2mp_bilateral MBONs should therefore lead to upregulation of *ban* target mRNAs. To identify candidate targets of *ban*, we performed fluorescence-activated cell sorting (FACS) and deep sequencing on γ5β′2a/β′2mp/β′2mp_bilateral MBONs expressing GFP and *ban-SP* or *scr-SP*. Adult brains were dissected at ZT12 (the time of maximum sleep-loss- Figure 1D). GFP+ MBONs were isolated by FACS and total mRNA was sequenced using a modified SMART-seq2 protocol (Liu et al., 2017; Picelli et al., 2014). RNA sequencing and Differential Gene Expression edgeR analysis (Robinson et al., 2010) for three biological replications was then performed to identify mRNAs that were significantly upregulated in the *ban* knockdown condition (Figure 2A). The list of statistically significant upregulated genes was compared to lists of putative target mRNAs generated by two microRNA-mRNA target prediction algorithms, TargetscanFly7.2 (Agarwal et al., 2018; Ruby et al., 2007) and DIANA-microT (Paraskevopoulou et al., 2013; Reczko et al., 2012). This pipeline produced 3 candidate mRNAs that were both significantly upregulated in the *ban* knockdown condition and were predicted direct targets-*UDP-glycosyltransferase family 36 member D1 (Ugt36D1)*, *Kelch (kel)* and *CCHamide-2 receptor (CCHa2-R)* (Figure 2B).

**Figure 2.**
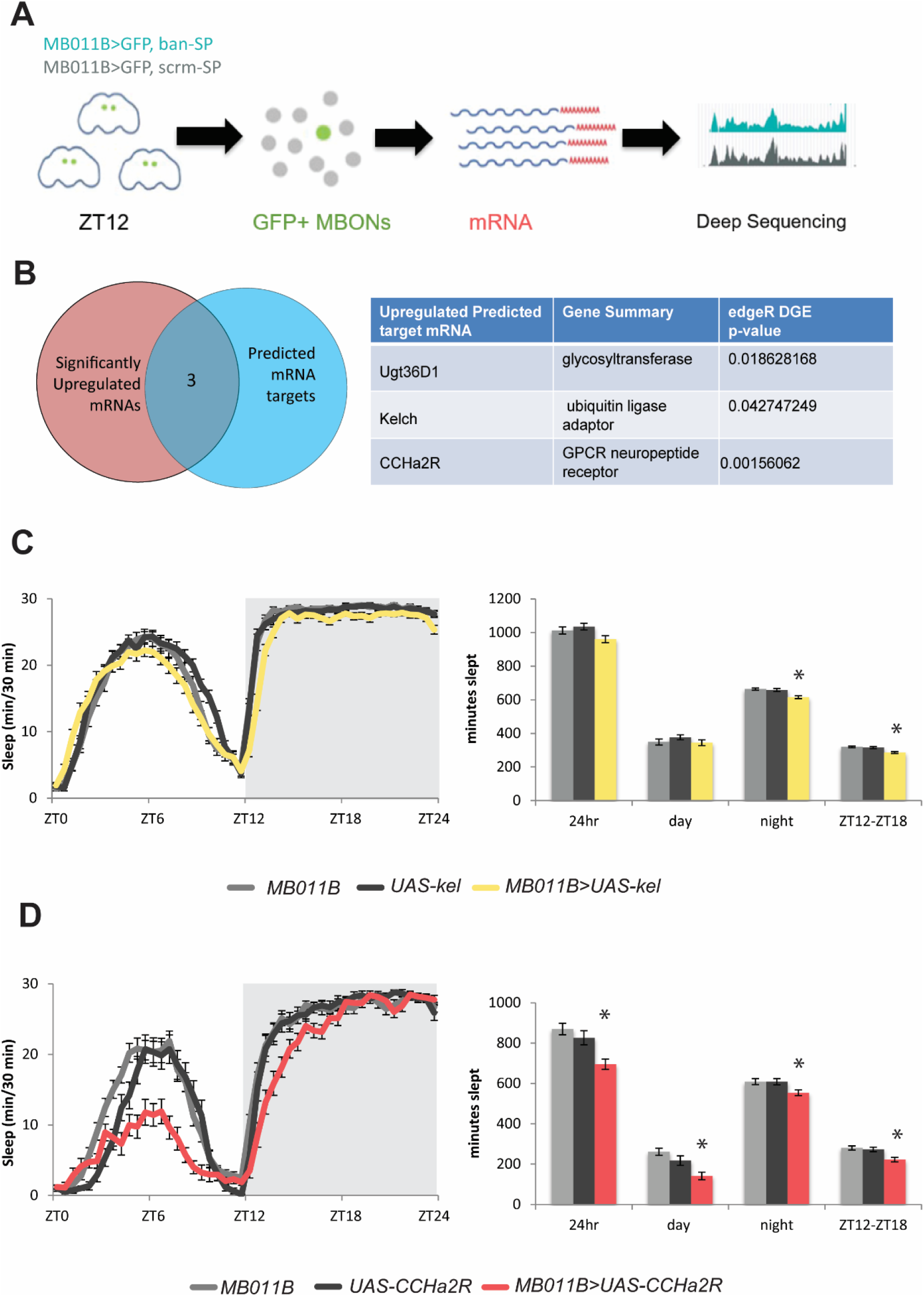
bantam negatively regulates wake-promoting mRNAs in the γ5β′2a/β′2mp/β′2mp_bilateral MBONs (A) Schematic diagram of MBON sequencing paradigm (B) Venn diagram representing selection criteria for putative ban microRNA targets in the γ5β′2a/β′2mp/β′2mp_bilateral MBONs. Table representing the three “hits”, their gene summarizations and their p-value significance for edgeR differential gene expression analysis. (C) Sleep data for *MB011B>UAS-kel* and parental line controls (mean± SEM). * represents P≤.001 using the Kruskal-Wallis Test with Dunn’s multiple comparison test. (D) Sleep data for *MB011B>UAS-CCHa2-R* and parental line controls (mean± SEM). * represents P≤.05 using the Kruskal-Wallis Test with Dunn’s multiple comparison test or one-way ANOVA with Tukey multiple comparisons test.

In order to determine if upregulation of these mRNAs is causative of the *ban* sleep loss phenotype, we overexpressed these genes within the γ5β′2a/β′2mp/β′2mp_bilateral MBONs and measured their effect on sleep. *MB011B>Ugt36D1* flies exhibited normal sleep (supplementary figure 3), but both *MB011B>kel* and *MB011B>CCHa2-R* flies exhibited statistically significant reductions in early nighttime sleep (Figure 2C and 2D), with overexpression of *kel* having only a modest effect. These results suggest that *ban* may mediate its nighttime sleep-promoting effect in the γ5β′2a/β′2mp/β′2mp_bilateral MBONs partially through the negative regulation of these target mRNAs.

Our results demonstrated that *ban* expression in the γ5β′2a/β′2mp/β′2mp_bilateral MBONs is required for normal early nighttime sleep. However, the underlying physiological processes regulated by *ban* within these neurons were unknown. Given the previously characterized role of *ban* in the regulation of cellular proliferation and differentiation (Banerjee and Roy, 2017; Brennecke et al., 2003; Lerner et al., 2015; Weng and Cohen, 2015), we asked whether knockdown of *ban* in γ5β′2a/β′2mp/β′2mp_bilateral MBONs led to an alteration in cell number or structure. Comparison of *MB011B>CD8::GFP+ ban-SP* to control *MB011B>CD8::GFP+scr-SP* brains revealed no differences in GFP+ cell number nor gross morphology (Supplementary Figure 4). These results suggest a novel adult role for *ban* in the regulation of behavior and cellular processes in the γ5β′2a/β′2mp/β′2mp_bilateral MBONs.

Given the normality of the structure of these MBONs, we hypothesized that *ban* had a role in regulation of their activity. It had been previously shown that *dTrpA1*-induced (Hamada et al., 2008) neural activation of the γ5β′2a/β′2mp/β′2mp_bilateral MBONs reduced sleep (Sitaraman et al., 2015a). In this same study, suppression of transmitter release using a temperature-sensitive dominant negative dynamin transgene (*shibire*^*ts*^) (Kitamoto, 2001), did not affect sleep, suggesting a model in which MBONs modify baseline sleep levels only in response to salient information presented to the MB. Given that neuronal activation of the γ5β′2a/β′2mp/β′2mp_bilateral MBONs phenocopied the effect of *ban* knockdown on sleep, this suggested that knockdown of *ban* may increase activity in these neurons.

To test this, we drove *GCaMP6f*, a calcium sensor, with *GMR14C08-GAL4,* a stronger driver for γ5β′2a/β′2mp/β′2mp_bilateral MBONs. MBONs are downstream of an excitatory pathway that conveys olfactory information to the MB circuit; antennal lobe (AL) projection neurons send excitatory inputs to Kenyon cells which in turn make excitatory connections upon MBONs (Aso et al., 2014a; Barnstedt et al., 2016; Ueno et al., 2013). This excitatory pathway was activated by a glass suction microelectrode placed on a subset of AL neurons and calcium dynamics in the dendrites of the ipsilateral group of γ5β′2a/β′2mp/β′2mp_bilateral MBONs were recorded (Figure 3A, B) (Ueno et al., 2013; Ueno et al., 2017; Wang et al., 2008). Consistent with our hypothesis, *GMR14C08>ban-SP+GCaMP6f* MBONs exhibited a significantly higher mean dendritic ΔF/F0 calcium signal than control *GMR14C08>scr+GCaMP6f* MBONs when the AL was stimulated at ZT12, the time of maximal sleep loss (Figures 3C and 3D). Since *ban* knockdown reduces sleep in a time-of-day-dependent manner we asked whether the effect of *ban* knockdown on activity exhibited a similar pattern. Mirroring our behavioral results, AL stimulation at ZT0 and ZT6 produced calcium responses which were statistically indistinguishable from *scr-SP* controls (Figure 3D). These results demonstrate that *ban* negatively regulates the activity of the γ5β′2a/β′2mp/β′2mp_bilateral MBONs during the early night, but not during the day, in line with its effects on sleep.

**Figure 3.**
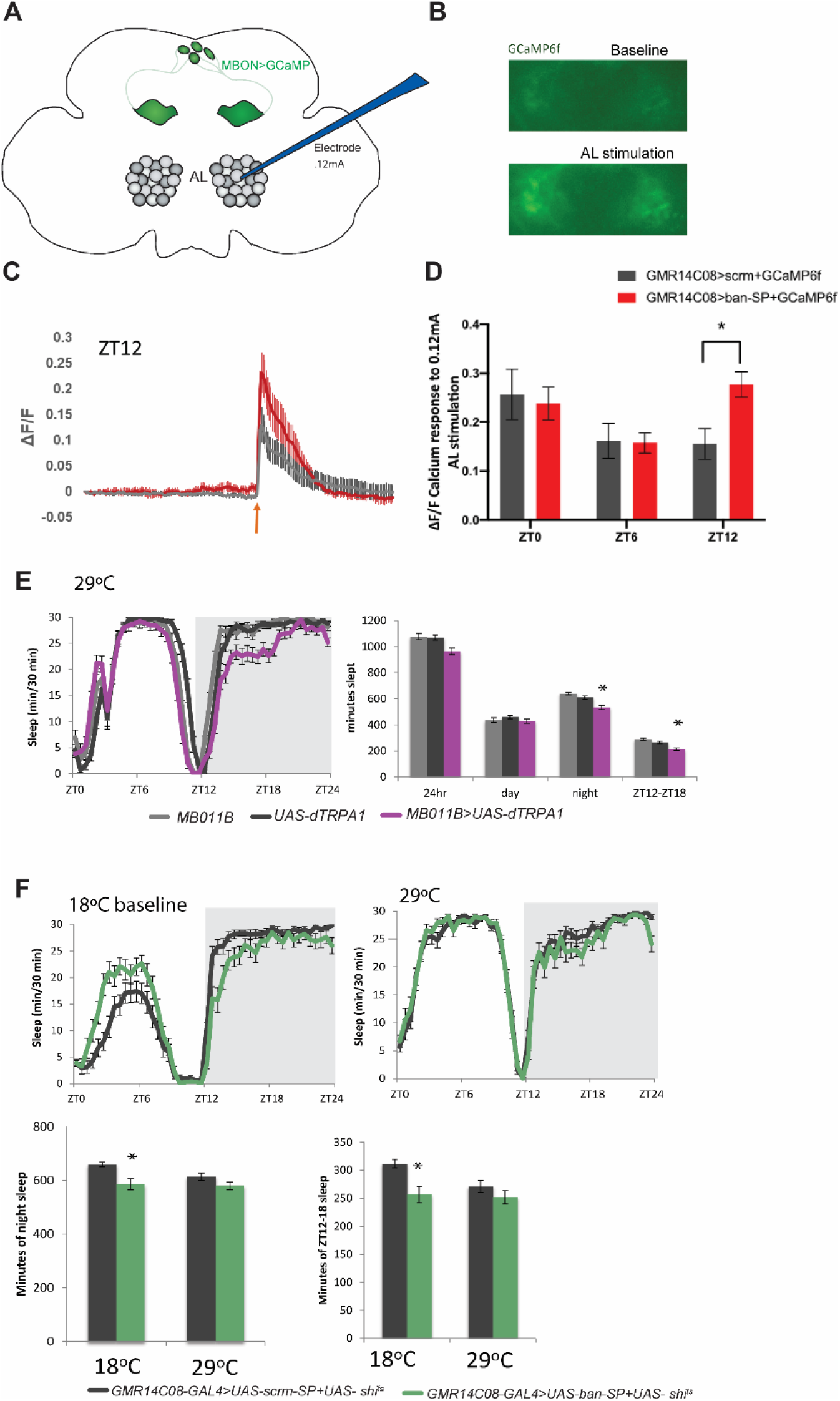
bantam inhibits neural activity in the wake-promoting γ5β′2a/β′2mp/β′2mp_bilateral MBONs (A) Schematic diagram of paradigm for antennal lobe (AL) microelectrode stimulation and γ5β′2a/β′2mp/β′2mp_bilateral MBON GCaMP6f recording (B) GCaMP6f images showing representative dendritic Ca^2+^ responses of *GMR14C08-GAL4>UAS-ban-SP+UAS-GCaMP6f* during baseline and during antennal lobe stimulation (C) ΔF/F time course for Ca^2+^ responses of *GMR14C08-GAL4>UAS-scrm-SP+UAS-GCaMP6f* (grey) and *GMR14C08-GAL4>UAS-ban-SP+UAS-GCaMP6f* (red) dendrites to antennal lobe stimulation (orange arrow) at ZT12 (mean± SEM). (D) ΔF/F for Ca^2+^ responses of *GMR14C08-GAL4>UAS-scrm-SP+UAS-GCaMP6f* (grey) and *GMR14C08-GAL4>UAS-ban-SP+UAS-GCaMP6f* (red) dendrites to antennal lobe stimulation (0.12mA) at ZT0, ZT6 and ZT12 (mean± SEM). * represents P≤.05 using two-way ANOVA and Sidak post hoc comparison. n of 8 for every timepoint. (E) Sleep data for *MB011B>UAS-dTrpA1* and parental line controls tested for sleep behavior at 29°C (mean± SEM). * represents P≤.01 using the Kruskal-Wallis Test with Dunn’s multiple comparison test. (F) Sleep data for *GMR14C08>UAS-ban-SP+UAS-shi*^*ts*^ and *GMR14C08>UAS-scrm-SP+UAS-shi*^*ts*^ tested at 18°C and 29°C (mean± SEM). * represents P≤.01 using the Mann-Whitney-Wilcoxon Test.

To ask if the increased neuronal activity associated with *ban* knockdown was causally related to the sleep loss phenotype we first confirmed that activation of the γ5β′2a/β′2mp/β′2mp_bilateral MBONs with the temperature-sensitive cation channel *dTrpA1* decreased nighttime sleep (Figure 3E) and that inhibition of MBON synaptic release with *shibire*^*ts*^ had no effect on baseline sleep (Supplementary Figure 5). We then hypothesized that if the hyperactivity associated with *ban* knockdown was causative for the sleep loss phenotype, inhibition of γ5β′2a/β′2mp/β′2mp_bilateral MBON activity in *ban-SP-*expressing flies should rescue early night sleep. To test this we co-expressed *ban-SP* with *shi*^*ts*^. *GMR14C08>ban-SP+UAS-shi*^*ts*^ flies exhibited decreased nighttime sleep when tested at the permissive temperature of 18°C (Figure 3F), consistent with loss of *ban* activity. However, when tested at the restrictive temperature of 30°C (which blocks MBON synaptic signaling), *GMR14C08-GAL4>ban-SP+shi*^*ts*^ flies exhibited sleep levels statistically indistinguishable from *scr-SP* controls (Figure 3F). The ability of *shi*^*ts*^ to rescue the sleep loss phenotype indicates that *ban* knockdown enhances neurotransmission by MBONs which in turn inhibits nighttime sleep. These results demonstrate that *ban*-mediated control of neuronal activity within the wake-promoting γ5β′2a/β′2mp/β′2mp_bilateral MBONs acts to promote early nighttime sleep.

## Discussion

microRNAs have emerged as key regulators of sleep and wake, but the cellular and physiological mechanisms via which individual microRNAs influence sleep remain largely uncharacterized. Here we show that the microRNA *bantam (ban)* promotes early nighttime sleep by decreasing the neuronal activity of the wake-promoting γ5β′2a, β′2mp, β′2mp_bilateral MBONs. *ban* is a widely-expressed microRNA with well-characterized developmental roles in the specification of neuron number and morphology (Parrish et al., 2009; Song et al., 2012; Weng and Cohen, 2015). Known adult roles for *ban* have been largely limited to regulation of stem cells or other proliferative populations (Huang et al., 2014; Shcherbata et al., 2007) making the adult-specific role of *ban* in regulation of neuronal excitability a novel function for this microRNA.

Down-regulation of *ban* produces a sleep loss phenotype largely restricted to the early night ZT12-18 period as well as an increase in sleep latency. A number of sleep-regulating genes exhibit patterns of sleep loss that are largely confined to the early night period (Cong et al., 2015) or have large effects on latency (Agosto et al., 2008). Conversely, other sleep-regulating genes primarily contribute to late night sleep (Gmeiner et al., 2013; Kunst et al., 2014). This may reflect the fact that early and late-night sleep represent physiologically distinct states under differential genetic and neuroanatomical regulation and which likely serve different functions. Mammalian sleep varies across the night with slow-wave non-REM sleep predominant during the early night and REM sleep more frequent during the late night. Early and late-night sleep have also been shown to facilitate different aspects of sleep-dependent memory processing (Plihal and Born, 1997; Yordanova et al., 2008). Our data show that *bantam* is important in allowing the initiation of early night sleep by suppressing activation of the γ5β′2a/β′2mp/β′2mp_bilateral MBONs.

Similar to mammalian sleep, *Drosophila* sleep is regulated by a widely distributed neural network (Shafer and Keene, 2021). The role of the mushroom body in the sculpting of responses to external conditions makes this structure uniquely suited to providing context-specific regulation of sleep. Consistent with this, synaptic silencing of the γ5β′2a/β′2mp/β′2mp_bilateral MBONs with *shi*^*ts*^ (Sitaraman et al., 2015a), electrical silencing (with *UAS-kir2.1*) and induced apoptosis (with *UAS-hid*) of the γ5β′2a/β′2mp/β′2mp_bilateral MBONs (data not shown) have no effect on basal sleep. A similar sleep regulation pattern was demonstrated for the wake-promoting *C01+A05* neurons: knockdown of the calcium sensor *Neurocalcin* in these cells led to reduced sleep and increased neural activity, whereas electrical silencing of the cells had no effect on sleep (Chen et al., 2019). This supports a framework in which many of the identified sleep-regulating neurons may actually be conditionally-recruited loci that allow specific internal and external states to influence immediate sleep/wake probability but do not have major effects on sleep in normal conditions.

We speculate that he γ5β′2a/β′2mp/β′2mp_bilateral MBONs are activated by physiological or environmental factors that reduce sleep in response to competing motivational drives, with *ban* expression acting as the switch to turn neural activity off and on. These MBON neurons are known to be required for behaviors that might compete with sleep, including avoidance of aversive stimuli, aversion to pathogen-infected food, ingestion, startle-induced locomotion, and memory for visual and olfactory cues (Al-Anzi and Zinn, 2018; Aso et al., 2014b; Kobler et al., 2020; Lewis et al., 2015; Owald et al., 2015; Sun et al., 2018; Yamazaki et al., 2018). For the γ5β′2a/β′2mp/β′2mp_bilateral MBONs, recruitment into the sleep circuitry likely also involves dopaminergic signaling since RNAi knockdown of the *DopR1* and *DopR2* dopamine receptors in these cells lead to an increase in nighttime sleep (Driscoll et al., 2020). Dopaminergic input to this subclass of MBONs was previously shown to be excitatory and wake-promoting (Sitaraman et al., 2015b). Thus dopamine signaling and *ban* expression have opposite effects on sleep and neural activity at this locus. Whether or not *ban* and dopamine act via modulation of the same pathways in the γ5β′2a/β′2mp/β′2mp_bilateral MBONs remains to be determined.

The sleep-promoting effect of *ban* in the γ5β′2a/β′2mp/β′2mp_bilateral MBONs appears to be the result of regulation of several mRNA targets, including *CCHamide-2 receptor* and *kelch*. *Kelch* is a BTB-domain adaptor protein for the Cullin-3 ubiquitin E3 ligase that is involved in actin regulation (Hudson and Cooley, 2010; Kelso et al., 2002) and dendritic branching (Djagaeva and Doronkin, 2009). Cullin-3 and *Insomniac*, another BTB-domain adaptor protein, have been previously implicated in sleep (Pfeiffenberger and Allada, 2012). *CCha2-R* is a GPCR for the gut-derived peptide hormone CCHamide-2 which has a critical role in feeding and growth (Ren et al., 2015; Sano et al., 2015). Activation of the CCHa2-R receptor enhances calcium responses in these neuroendocrine cells (Sano et al., 2015), consistent with a role for CCHa2-R in promoting neural activity of MBONs. Activation of BRS3 (Bombesin receptor subtype-3), the human ortholog of CCHa2-R depolarizes sleep-regulating orexin neurons in mice (Furutani et al., 2010), implying a parallel function for this receptor in activity regulation of mammalian sleep-relevant neurons. Interestingly, the *BRS3* mRNA hosts an MRE for the mammalian ortholog of *ban*, miR-450b-3p (Ibanez-Ventoso et al., 2008; Paraskevopoulou et al., 2013; Reczko et al., 2012) suggesting a highly conserved functional interaction between *ban* and this class of receptor.

## Methods

### *Drosophila* Lines

UAS lines include *UAS-ban-SP* and *UAS-scramble-SP* (Fulga et al., 2015), *UAS-CD8::GFP* (Lee and Luo, 1999), *UAS-kelch* (Hudson et al., 2015), *20XUAS-GCaMP6f* (Chen et al., 2013), *20XUAS-IVS-dTrpA1* (Hamada et al., 2008) and *20xUAS-IVS-Syn21-Shi*^*ts*^ (Pfeiffer et al., 2012). *UAS-Ugt36D1* and *UAS-CCha2-R* lines were generated by cloning their cDNA sequences into the *Drosophila e*xpression vector *pUASTattB* which was then integrated into the attp40 PhiC31 integration site on chromosome 2 (Bischof et al., 2007). Injections were performed by Rainbow Transgenic Flies Inc (Camarillo, CA).GAL4s include *nSyb-GAL4*, *ban-Gal4* (generous gift of Sebastian Kadener, Brandeis University), *OK107-GAL4* (Aso et al., 2009), *TH-GAL4* (Friggi-Grelin et al., 2003) and *GMR14C08-GAL4* (Jenett et al., 2012). Split-GAL4s include *MB011B* (Aso et al., 2014a). GAL80 lines include *tubulin-GAL80*^*ts*^ (McGuire et al., 2003).

### *Drosophila* Husbandry and Sleep Assay

Flies were grown on standard cornmeal/ agar food supplemented with yeast. *Drosophila* Activity Monitor (DAM) system (TriKinetics, Waltham) was used to measure sleep (Donelson et al., 2012). Female flies were loaded into glass sleep tubes containing a food mixture of 5% sucrose and 2% agar. Young female flies were housed with W^1118^ males >24 hours prior to loading in sleep tubes in order to ensure that flies were mated. Temperature was kept constant at 25°C throughout sleep recording unless otherwise noted. Flies were considered sleeping if they were inactive for five or more minutes (Hendricks et al., 2000; Shaw et al., 2000). Sleep was averaged across 2-5 days unless otherwise noted. DAM data was analyzed using a custom MATLAB program called Sleep and Circadian Analysis MATLAB Program (SCAMP)(Donelson et al., 2012). For all sleep duration data, a D’Agostino-Pearson test was used to test for normality. If normally distributed, data was tested with ANOVA or t test depending on the number of groups. If not normally distributed, data was tested with a Kruskal-Wallis ANOVA with Dunn’s multiple comparison test or with a Mann-Whitney test. Statistics were performed using GraphPad Prism.

### Immunohistochemistry

Fly brains were dissected in ice cold Schneider’s Insect Medium (S2) and fixed in 2% paraformaldehyde for 55 minutes at room temperature. Brains were then washed 4X in PBS with .5% Triton X-100 (PBS-T) and then placed in blocking solution (PBS-T with 5% normal goat serum (NGS; Invitrogen)) for 90 minutes at room temperature. Incubation in primary and secondary antibodies was performed for 2-3 days at 4°C. Anti-GFP (raised in rabbit; Invitrogen) and anti-BRP (raised in mouse; monoclonal Nc82) were used at concentrations 1:1000 and 1:25, respectively. Anti-rabbit Alexa Flour 488 and anti-mouse Alexa Flour 633 (both Invitrogen) were used at 1:500. Brains were then washed 4X times in PBS-T and placed in 2% paraformaldehyde for 4 hours at room temperature. Brains were washed 4X times in PBS-T and then mounted using Vectashield Mounting Medium (Vector Laboratories). Slides were imaged on a Leica SP5 confocal microscope with a 63x objective. Maximum intensity Z projections were generated using FIJI software.

### FACS Sorting and RNA-seq of MBONs

FACS was performed using a previously described protocol (Ma et al., 2021). The brains of *MB011B>UAS-CD8::GFP+UAS-ban-SP* and control *MB011B>UAS-CD8::GFP+UAS-scrm-SP* mated female flies (3-4 days post-eclosion) were dissected in artificial hemolymph (AHL-108 mM NaCl, 5 mM KCl, 2 mM CaCl2, 8.2 mM MgCl2, 4 mM NaHCO3, 1 mM NaH2PO4-H2O, 5 mM trehalose, 10 mM sucrose, 5 mM HEPES; pH 7.5) and 0.1 μM tetrodotoxin (TTX). Brains were collected in SM medium (active Schneider’s medium) and 0.1 μM TTX. Brains were then digested in Papain (Worthington PAP2, 50 unit/ml, with approximately 2 μl per brain) for 30 min and then quenched with 5X volume of SM media+TTX. Trituration was performed with flame-rounded 1000 μl and 200 μl pipette tips. Filtration was performed with a 100 μm sieve. Cell sorting was performed using a BD FACS Melody using the GFP channel. Sorted cells were collected in an Eppendorf tube with 50μl lysis buffer (Dynabead mRNA purification kit, Invitrogen) and stored at −80°C.

RNA sequencing was performed using a modified SMART-seq2 protocol (Li et al., 2017; Picelli et al., 2014). Briefly, mRNA was isolated from cells using a Dynabead mRNA purification kit (Invitrogen). Poly(A)-tailed RNA was reverse transcribed and PCR-amplified with 25 cycles. cDNA libraries were cleaned using AMPure beads (Agencourt). The resulting full-length cDNA was used as the input to the tagmentation based Nextera XT (Illumina) protocol to generate sequencing libraries. Two biological replications were sequenced on a Next-seq 500 platform with paired-end 75bp reads and one biological replication was sequenced on a Hi-seq platform with paired-end 150bp reads. Reads were aligned to the *Drosophila* genome (dm6) using STAR (Dobin et al., 2013). EdgeR was used to perform DGE on the aligned reads of the three biological replicates (Robinson et al., 2010). The list of predicted *ban* targets was generated by TargetscanFly7.2 (Agarwal et al., 2018; Ruby et al., 2007) and DIANA-microT (Paraskevopoulou et al., 2013; Reczko et al., 2012) using an miTG threshold of .5.

### GCaMP6f Imaging and Antennal Lobe Stimulation

Calcium imaging was performed using a modified version of a previously published protocol (Ueno et al., 2013). All imaged flies were mated females, 4-5 days post-eclosion. *GMR14C08 >UAS-ban-SP+GCaMP6f* and *GMR14C08 >UAS-scrm-SP+GCaMP6f* brains were dissected in ice cold HL3.1 (Feng et al., 2004) and loaded into a recording chamber submerged in HL3.1. A glass microelectrode was used to stimulate one antennal lobe. The microelectrode was approximately one fourth the size of the antennal lobe. The microelectrode delivered a stimulation train of 20 0.12mA pulses at 100hz with a pulse width of 1 millisecond and an inter-pulse interval of 9 milliseconds.

Simultaneously, GCaMP signals were recorded from the dendritic arborizations of the β′2a/β′2mp/β′2mp_bilateral MBONs (identifiable by their distinct morphology). Imaging data was collected at an acquisition rate of 10Hz (100 millisecond exposure) at a 512×512 resolution using an Olympus microscope with a 40x objective. Only MBON dendrites ipsilateral to the stimulated Antennal Lobe were included in analysis. A ΔF/F0 was calculated for the baseline period (5 seconds prior to stimulus onset) and the max calcium response during antennal lobe stimulation. Statistical analysis (two-way ANOVA with Sidak post-hoc test) was performed using GraphPad Prism.

---

**Supplementary Figure 1.**
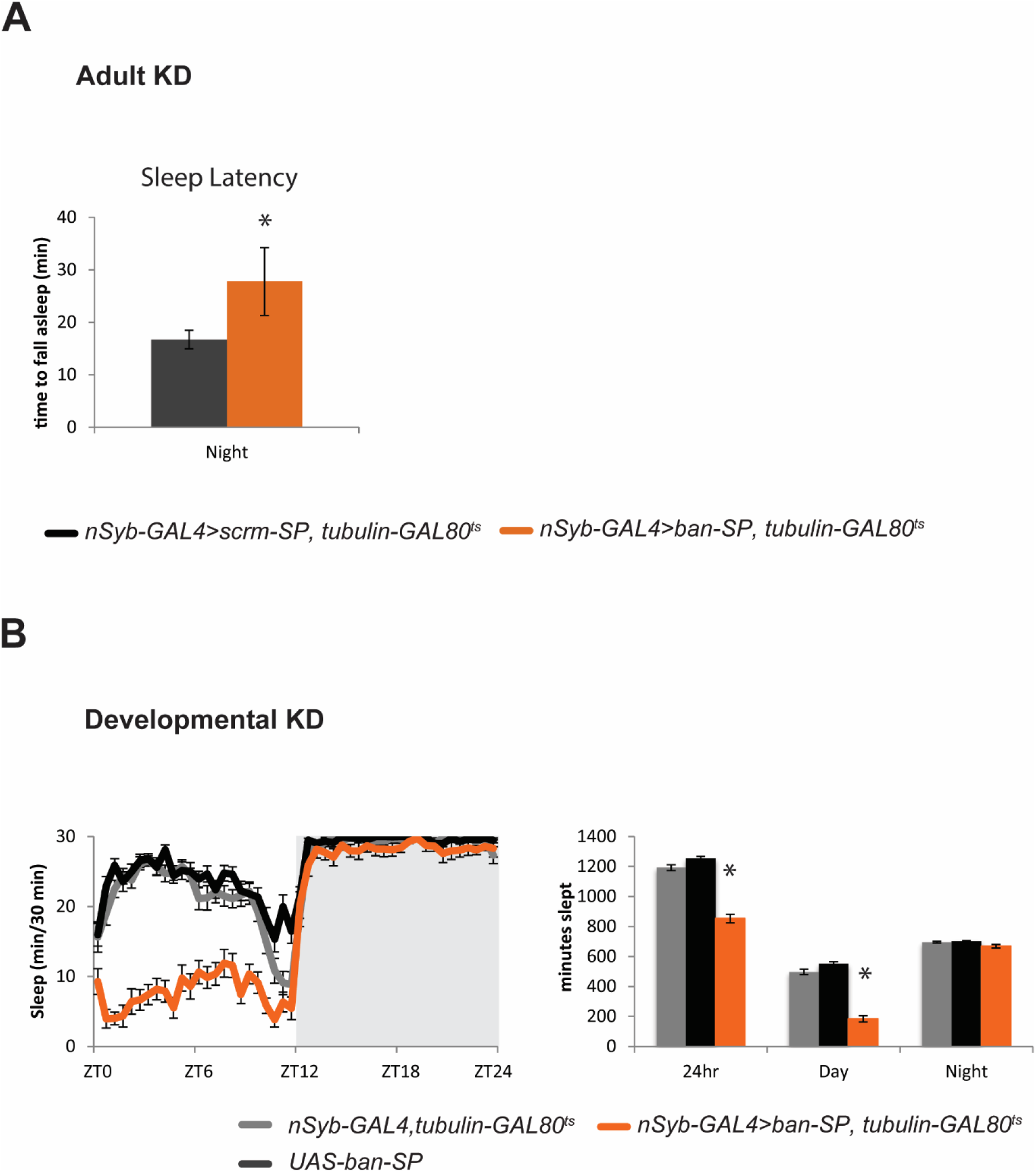
(A) Sleep latency (minutes to fall asleep after night onset) for *nSyb-GAL4>UAS-ban-SP, tubulin-GAL80*^*ts*^ and scramble controls raised at 17°C and tested at 29°C (mean± SEM). * represents P≤.0001 for an unpaired t test. (B) Developmental knockdown of *ban*. Sleep data for *nSyb-GAL4>UAS-ban-SP, tubulin-GAL80*^*ts*^ and parental line controls raised at 29°C and tested at 18°C (mean± SEM). * represents P≤.0001 using the Kruskal-Wallis Test with Dunn’s multiple comparison test.

**Supplementary Figure 2.**
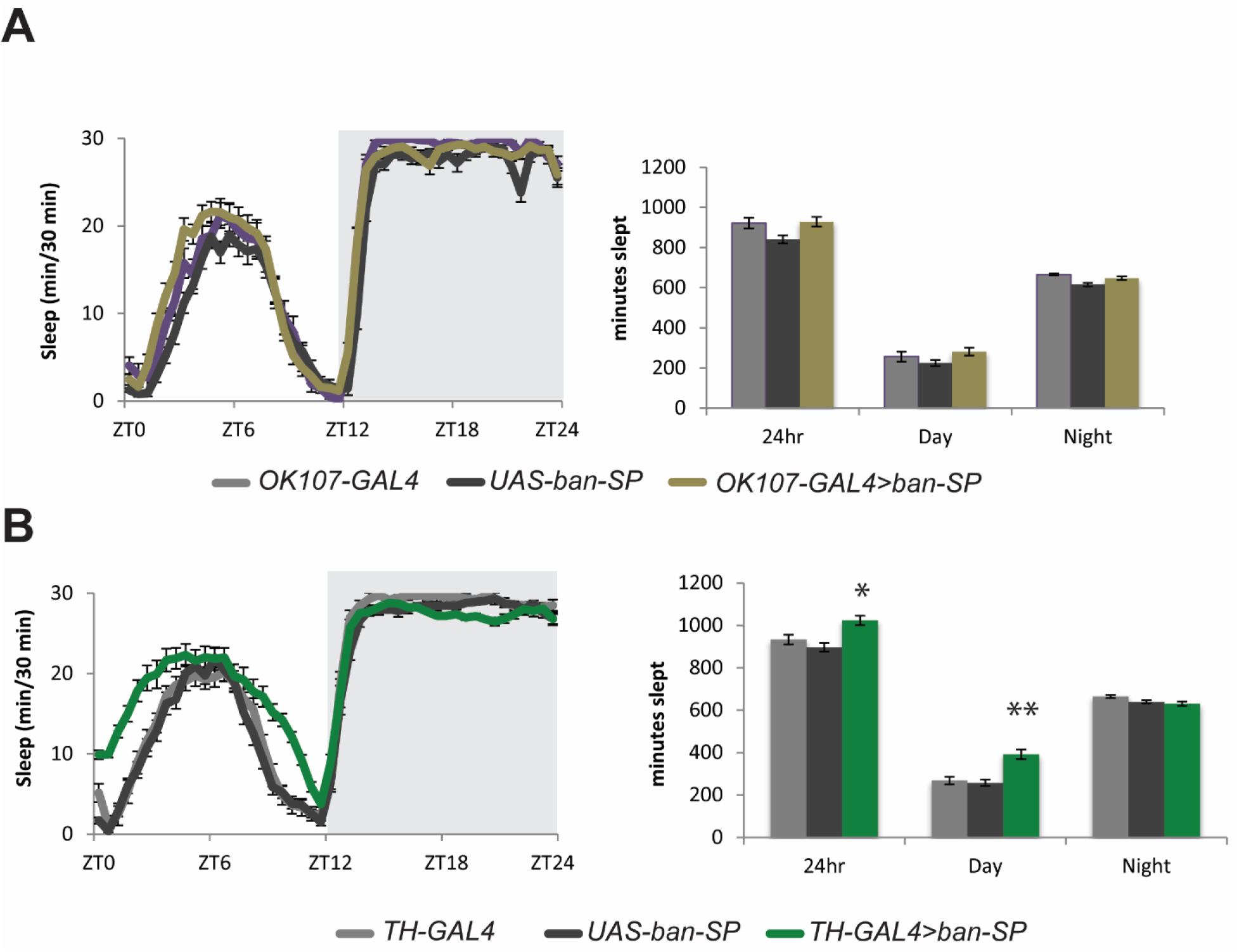
(A) Sleep data for *OK107-GAL4>UAS-ban-SP* and parental line controls (mean± SEM). (B) Sleep data for *TH-GAL4>UAS-ban-SP* and parental line controls (mean± SEM). * represents P≤.05 using the Kruskal-Wallis Test with Dunn’s multiple comparison test. ** represents P≤.0001 using a one-way ANOVA with Tukey’s multiple comparison test.

**Supplementary Figure 3.**
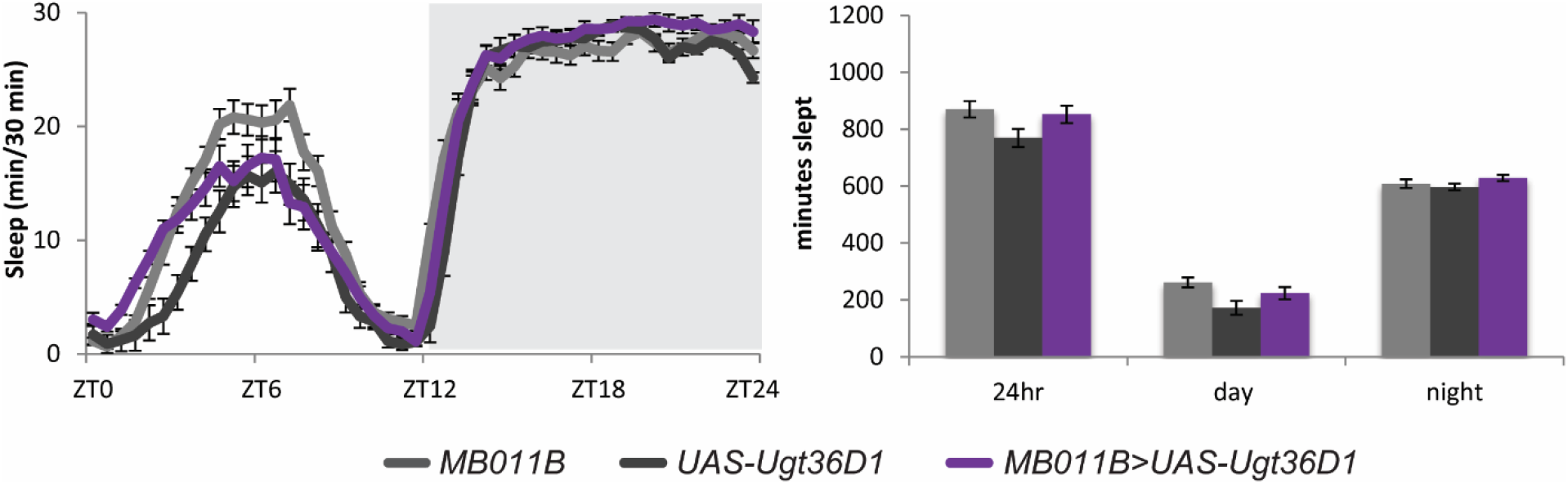
(A) Sleep data for *MB011B>UAS-Ugt36D1* and parental line controls (mean± SEM).

**Supplementary Figure 4.**
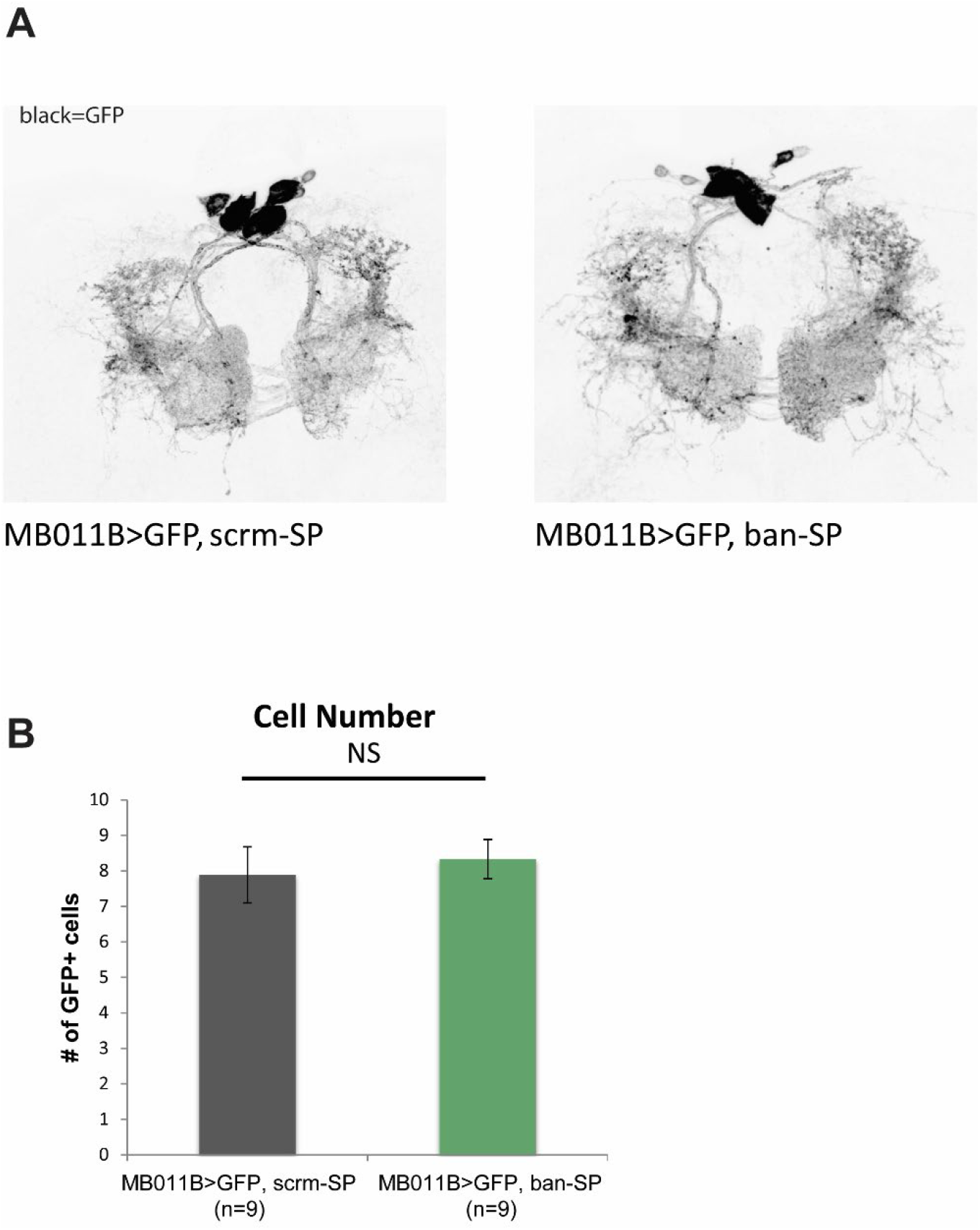
(A) IHC anti-GFP staining for representative central brains of *MB011B>UAS-CD8::GFP+UAS-scrm-SP* and *MB011B>UAS-CD8::GFP+UAS-ban-SP* flies. (B) Quantification of the number of GFP+ cells per brain.

**Supplementary Figure 5.**
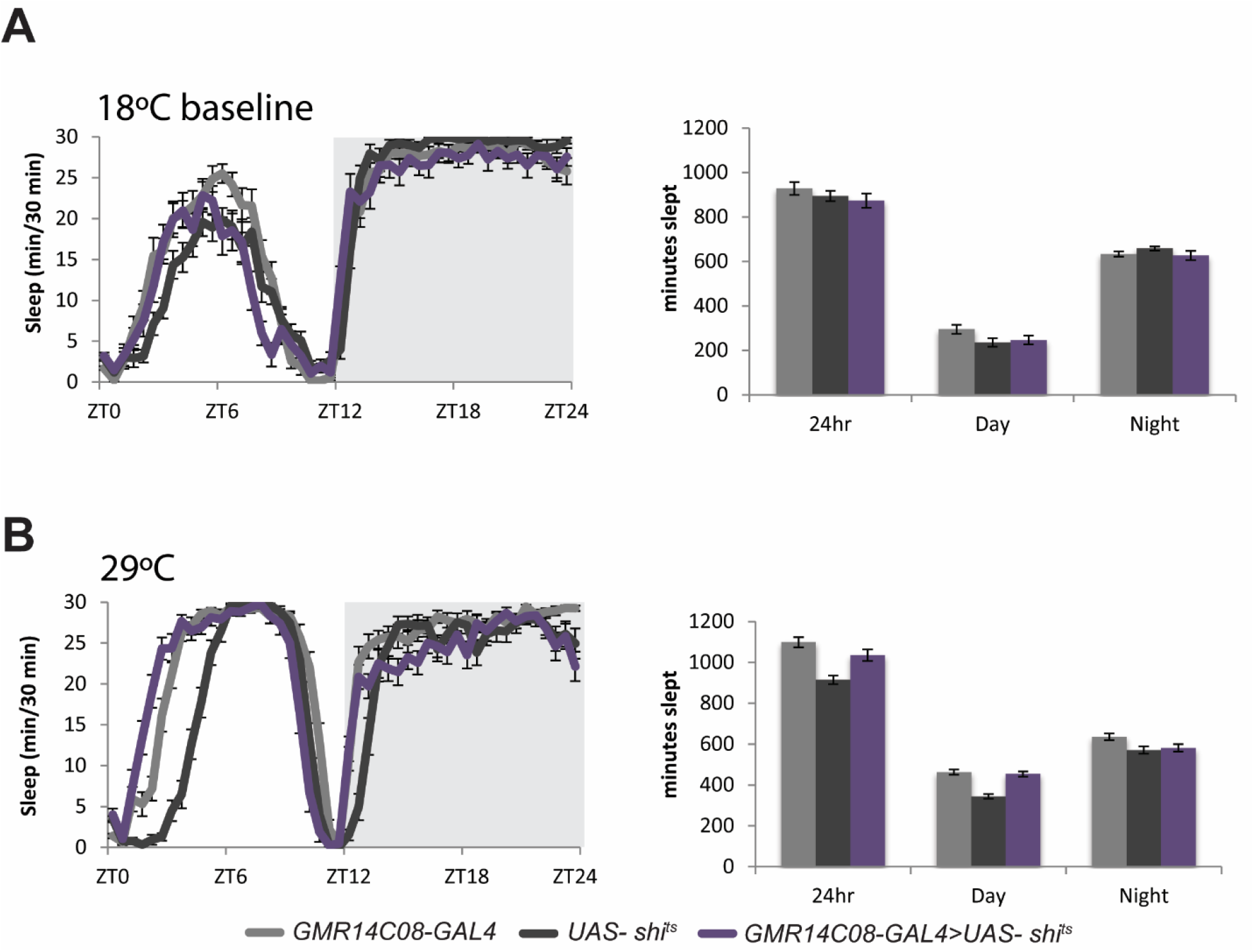
(A) Sleep data for *GMR14C08-GAL4>UAS-shi*^*ts*^ and parental line controls at 18°C (mean± SEM). (B) Sleep data for *GMR14C08-GAL4>UAS-shi*^*ts*^ and parental line controls at 29°C (mean± SEM).

## References

Agarwal, V., Subtelny, A.O., Thiru, P., Ulitsky, I., and Bartel, D.P. (2018). Predicting microRNA targeting efficacy in Drosophila. Genome biology 19, 152.

Agosto, J., Choi, J.C., Parisky, K.M., Stilwell, G., Rosbash, M., and Griffith, L.C. (2008). Modulation of GABA_A_ receptor desensitization uncouples sleep onset and maintenance in *Drosophila*. Nat Neurosci 11, 354–359.

Al-Anzi, B., and Zinn, K. (2018). Identification and characterization of mushroom body neurons that regulate fat storage in Drosophila. Neural Dev 13, 18.

Aso, Y., Grubel, K., Busch, S., Friedrich, A.B., Siwanowicz, I., and Tanimoto, H. (2009). The mushroom body of adult Drosophila characterized by GAL4 drivers. J Neurogenet 23, 156–172.

Aso, Y., Hattori, D., Yu, Y., Johnston, R.M., Iyer, N.A., Ngo, T.T., Dionne, H., Abbott, L.F., Axel, R., Tanimoto, H., et al. (2014a). The neuronal architecture of the mushroom body provides a logic for associative learning. Elife 3, e04577.

Aso, Y., Sitaraman, D., Ichinose, T., Kaun, K.R., Vogt, K., Belliart-Guerin, G., Placais, P.Y., Robie, A.A., Yamagata, N., Schnaitmann, C., et al. (2014b). Mushroom body output neurons encode valence and guide memory-based action selection in Drosophila. Elife 3, e04580.

Banerjee, A., and Roy, J.K. (2017). Dicer-1 regulates proliferative potential of Drosophila larval neural stem cells through bantam miRNA based down-regulation of the G1/S inhibitor Dacapo. Dev Biol 423, 57–65.

Barnstedt, O., Owald, D., Felsenberg, J., Brain, R., Moszynski, J.P., Talbot, C.B., Perrat, P.N., and Waddell, S. (2016). Memory-Relevant Mushroom Body Output Synapses Are Cholinergic. Neuron 89, 1237–1247.

Beckwith, E.J., and French, A.S. (2019). Sleep in Drosophila and Its Context. Front Physiol 10, 1167.

Bischof, J., Maeda, R.K., Hediger, M., Karch, F., and Basler, K. (2007). An optimized transgenesis system for Drosophila using germ-line-specific phiC31 integrases. Proc Natl Acad Sci U S A 104, 3312–3317.

Brennecke, J., Hipfner, D.R., Stark, A., Russell, R.B., and Cohen, S.M. (2003). bantam encodes a developmentally regulated microRNA that controls cell proliferation and regulates the proapoptotic gene hid in Drosophila. Cell 113, 25–36.

Chakravarti Dilley, L., Szuperak, M., Gong, N.N., Williams, C.E., Saldana, R.L., Garbe, D.S., Syed, M.H., Jain, R., and Kayser, M.S. (2020). Identification of a molecular basis for the juvenile sleep state. Elife 9.

Chen, K.F., Lowe, S., Lamaze, A., Kratschmer, P., and Jepson, J. (2019). Neurocalcin regulates nighttime sleep and arousal in Drosophila. Elife 8.

Chen, T.W., Wardill, T.J., Sun, Y., Pulver, S.R., Renninger, S.L., Baohan, A., Schreiter, E.R., Kerr, R.A., Orger, M.B., Jayaraman, V., et al. (2013). Ultrasensitive fluorescent proteins for imaging neuronal activity. Nature 499, 295–300.

Chen, X., and Rosbash, M. (2017). MicroRNA-92a is a circadian modulator of neuronal excitability in Drosophila. Nat Commun 8, 14707.

Cong, X., Wang, H., Liu, Z., He, C., An, C., and Zhao, Z. (2015). Regulation of Sleep by Insulin-like Peptide System in Drosophila melanogaster. Sleep 38, 1075–1083.

Crocker, A., and Sehgal, A. (2010). Genetic analysis of sleep. Genes Dev 24, 1220–1235.

Davis, C.J., Bohnet, S.G., Meyerson, J.M., and Krueger, J.M. (2007). Sleep loss changes microRNA levels in the brain: a possible mechanism for state-dependent translational regulation. Neurosci Lett 422, 68–73.

Davis, C.J., Clinton, J.M., and Krueger, J.M. (2012). MicroRNA 138, let-7b, and 125a inhibitors differentially alter sleep and EEG delta-wave activity in rats. J Appl Physiol (1985) 113, 1756–1762.

Davis, C.J., Clinton, J.M., Taishi, P., Bohnet, S.G., Honn, K.A., and Krueger, J.M. (2011). MicroRNA 132 alters sleep and varies with time in brain. J Appl Physiol (1985) 111, 665–672.

Djagaeva, I., and Doronkin, S. (2009). COP9 limits dendritic branching via Cullin3-dependent degradation of the actin-crosslinking BTB-domain protein Kelch. PLoS One 4, e7598.

Dobin, A., Davis, C.A., Schlesinger, F., Drenkow, J., Zaleski, C., Jha, S., Batut, P., Chaisson, M., and Gingeras, T.R. (2013). STAR: ultrafast universal RNA-seq aligner. Bioinformatics 29, 15–21.

Donelson, N.C., Kim, E.Z., Slawson, J.B., Vecsey, C.G., Huber, R., and Griffith, L.C. (2012). High-resolution positional tracking for long-term analysis of Drosophila sleep and locomotion using the “tracker” program. PLoS One 7, e37250.

Driscoll M, Victoria C, Morgan M, Amanda N, Divya S. 2020. Dopamine neurons promote arousal and wakefulness via dop1R receptor in the drosophila mushroom body. BioRxiv. May, 2020.04.29.069229 (https://doi.org/10.1101/2020.04.29.069229, accessed August 21st, 2021).

Feng, Y., Ueda, A., and Wu, C.F. (2004). A modified minimal hemolymph-like solution, HL3.1, for physiological recordings at the neuromuscular junctions of normal and mutant Drosophila larvae. J Neurogenet 18, 377–402.

Friggi-Grelin, F., Coulom, H., Meller, M., Gomez, D., Hirsh, J., and Birman, S. (2003). Targeted gene expression in Drosophila dopaminergic cells using regulatory sequences from tyrosine hydroxylase. J Neurobiol 54, 618–627.

Fulga, T.A., McNeill, E.M., Binari, R., Yelick, J., Blanche, A., Booker, M., Steinkraus, B.R., Schnall-Levin, M., Zhao, Y., DeLuca, T., et al. (2015). A transgenic resource for conditional competitive inhibition of conserved Drosophila microRNAs. Nature communications 6, 7279.

Furutani, N., Hondo, M., Tsujino, N., and Sakurai, T. (2010). Activation of bombesin receptor subtype-3 influences activity of orexin neurons by both direct and indirect pathways. J Mol Neurosci 42, 106–111.

Gmeiner, F., Kolodziejczyk, A., Yoshii, T., Rieger, D., Nassel, D.R., and Helfrich-Forster, C. (2013). GABA(B) receptors play an essential role in maintaining sleep during the second half of the night in Drosophila melanogaster. J Exp Biol 216, 3837–3843.

Gong, Z., Xia, S., Liu, L., Feng, C., and Guo, A. (1998). Operant visual learning and memory in Drosophila mutants dunce, amnesiac and radish. J Insect Physiol 44, 1149–1158.

Goodwin, P.R., Meng, A., Moore, J., Hobin, M., Fulga, T.A., Van Vactor, D., and Griffith, L.C. (2018). MicroRNAs Regulate Sleep and Sleep Homeostasis in Drosophila. Cell reports 23, 3776–3786.

Hamada, F.N., Rosenzweig, M., Kang, K., Pulver, S.R., Ghezzi, A., Jegla, T.J., and Garrity, P.A. (2008). An internal thermal sensor controlling temperature preference in Drosophila. Nature 454, 217–220.

Hendricks, J.C., Finn, S.M., Panckeri, K.A., Chavkin, J., Williams, J.A., Sehgal, A., and Pack, A.I. (2000). Rest in Drosophila is a sleep-like state. Neuron 25, 129–138.

Holm, A., Bang-Berthelsen, C.H., Knudsen, S., Kornum, B.R., Modvig, S., Jennum, P., and Gammeltoft, S. (2014). miRNA profiles in plasma from patients with sleep disorders reveal dysregulation of miRNAs in narcolepsy and other central hypersomnias. Sleep 37, 1525–1533.

Huang, X., Shi, L., Cao, J., He, F., Li, R., Zhang, Y., Miao, S., Jin, L., Qu, J., Li, Z., et al. (2014). The sterile 20-like kinase tao controls tissue homeostasis by regulating the hippo pathway in Drosophila adult midgut. J Genet Genomics 41, 429–438.

Hudson, A.M., and Cooley, L. (2010). Drosophila Kelch functions with Cullin-3 to organize the ring canal actin cytoskeleton. J Cell Biol 188, 29–37.

Hudson, A.M., Mannix, K.M., and Cooley, L. (2015). Actin Cytoskeletal Organization in Drosophila Germline Ring Canals Depends on Kelch Function in a Cullin-RING E3 Ligase. Genetics 201, 1117–1131.

Huntzinger, E., and Izaurralde, E. (2011). Gene silencing by microRNAs: contributions of translational repression and mRNA decay. Nat Rev Genet 12, 99–110.

Ibanez-Ventoso, C., Vora, M., and Driscoll, M. (2008). Sequence relationships among C. elegans, D. melanogaster and human microRNAs highlight the extensive conservation of microRNAs in biology. PLoS One 3, e2818.

Jenett, A., Rubin, G.M., Ngo, T.T., Shepherd, D., Murphy, C., Dionne, H., Pfeiffer, B.D., Cavallaro, A., Hall, D., Jeter, J., et al. (2012). A GAL4-driver line resource for Drosophila neurobiology. Cell Rep 2, 991–1001.

Jiang, N., Soba, P., Parker, E., Kim, C.C., and Parrish, J.Z. (2014). The microRNA bantam regulates a developmental transition in epithelial cells that restricts sensory dendrite growth. Development 141, 2657–2668.

Joiner, W.J., Crocker, A., White, B.H., and Sehgal, A. (2006). Sleep in Drosophila is regulated by adult mushroom bodies. Nature 441, 757–760.

Jonas, S., and Izaurralde, E. (2015). Towards a molecular understanding of microRNA-mediated gene silencing. Nat Rev Genet 16, 421–433.

Kadener, S., Menet, J.S., Sugino, K., Horwich, M.D., Weissbein, U., Nawathean, P., Vagin, V.V., Zamore, P.D., Nelson, S.B., and Rosbash, M. (2009). A role for microRNAs in the Drosophila circadian clock. Genes Dev 23, 2179–2191.

Kelso, R.J., Hudson, A.M., and Cooley, L. (2002). Drosophila Kelch regulates actin organization via Src64-dependent tyrosine phosphorylation. J Cell Biol 156, 703–713.

Kitamoto, T. (2001). Conditional modification of behavior in Drosophila by targeted expression of a temperature-sensitive shibire allele in defined neurons. J Neurobiol 47, 81–92.

Kobler, J.M., Rodriguez Jimenez, F.J., Petcu, I., and Grunwald Kadow, I.C. (2020). Immune Receptor Signaling and the Mushroom Body Mediate Post-ingestion Pathogen Avoidance. Curr Biol 30, 4693–4709 e4693.

Kunst, M., Hughes, M.E., Raccuglia, D., Felix, M., Li, M., Barnett, G., Duah, J., and Nitabach, M.N. (2014). Calcitonin gene-related peptide neurons mediate sleep-specific circadian output in Drosophila. Curr Biol 24, 2652–2664.

Lee, T., and Luo, L. (1999). Mosaic analysis with a repressible cell marker for studies of gene function in neuronal morphogenesis. Neuron 22, 451–461.

Lerner, I., Bartok, O., Wolfson, V., Menet, J.S., Weissbein, U., Afik, S., Haimovich, D., Gafni, C., Friedman, N., Rosbash, M., et al. (2015). Clk post-transcriptional control denoises circadian transcription both temporally and spatially. Nature communications 6, 7056.

Lewis, L.P., Siju, K.P., Aso, Y., Friedrich, A.B., Bulteel, A.J., Rubin, G.M., and Grunwald Kadow, I.C. (2015). A Higher Brain Circuit for Immediate Integration of Conflicting Sensory Information in Drosophila. Curr Biol 25, 2203–2214.

Li, H., Horns, F., Wu, B., Xie, Q., Li, J., Li, T., Luginbuhl, D.J., Quake, S.R., and Luo, L. (2017). Classifying Drosophila Olfactory Projection Neuron Subtypes by Single-Cell RNA Sequencing. Cell 171, 1206–1220 e1222.

Liu, H., Li, Y., He, J., Guan, Q., Chen, R., Yan, H., Zheng, W., Song, K., Cai, H., Guo, Y., et al. (2017). Robust transcriptional signatures for low-input RNA samples based on relative expression orderings. BMC Genomics 18, 913.

Loya, C.M., Lu, C.S., Van Vactor, D., and Fulga, T.A. (2009). Transgenic microRNA inhibition with spatiotemporal specificity in intact organisms. Nature methods 6, 897–903.

Ma, D., Przybylski, D., Abruzzi, K.C., Schlichting, M., Li, Q., Long, X., and Rosbash, M. (2021). A transcriptomic taxonomy of Drosophila circadian neurons around the clock. Elife 10.

Mao, Z., and Davis, R.L. (2009). Eight different types of dopaminergic neurons innervate the Drosophila mushroom body neuropil: anatomical and physiological heterogeneity. Frontiers in neural circuits 3, 5.

McGuire, S.E., Le, P.T., Osborn, A.J., Matsumoto, K., and Davis, R.L. (2003). Spatiotemporal rescue of memory dysfunction in Drosophila. Science 302, 1765–1768.

Nitz, D.A., van Swinderen, B., Tononi, G., and Greenspan, R.J. (2002). Electrophysiological correlates of rest and activity in Drosophila melanogaster. Curr Biol 12, 1934–1940.

Owald, D., Felsenberg, J., Talbot, C.B., Das, G., Perisse, E., Huetteroth, W., and Waddell, S. (2015). Activity of defined mushroom body output neurons underlies learned olfactory behavior in Drosophila. Neuron 86, 417–427.

Paraskevopoulou, M.D., Georgakilas, G., Kostoulas, N., Vlachos, I.S., Vergoulis, T., Reczko, M., Filippidis, C., Dalamagas, T., and Hatzigeorgiou, A.G. (2013). DIANA-microT web server v5.0: service integration into miRNA functional analysis workflows. Nucleic Acids Res 41, W169–173.

Parrish, J.Z., Xu, P., Kim, C.C., Jan, L.Y., and Jan, Y.N. (2009). The microRNA bantam functions in epithelial cells to regulate scaling growth of dendrite arbors in drosophila sensory neurons. Neuron 63, 788–802.

Pfeiffenberger, C., and Allada, R. (2012). Cul3 and the BTB adaptor insomniac are key regulators of sleep homeostasis and a dopamine arousal pathway in Drosophila. PLoS Genet 8, e1003003.

Pfeiffer, B.D., Truman, J.W., and Rubin, G.M. (2012). Using translational enhancers to increase transgene expression in Drosophila. Proc Natl Acad Sci U S A 109, 6626–6631.

Picelli, S., Faridani, O.R., Bjorklund, A.K., Winberg, G., Sagasser, S., and Sandberg, R. (2014). Full-length RNA-seq from single cells using Smart-seq2. Nat Protoc 9, 171–181.

Pitman, J.L., McGill, J.J., Keegan, K.P., and Allada, R. (2006). A dynamic role for the mushroom bodies in promoting sleep in *Drosophila*. Nature 441, 753–756.

Plihal, W., and Born, J. (1997). Effects of early and late nocturnal sleep on declarative and procedural memory. J Cogn Neurosci 9, 534–547.

Reczko, M., Maragkakis, M., Alexiou, P., Grosse, I., and Hatzigeorgiou, A.G. (2012). Functional microRNA targets in protein coding sequences. Bioinformatics 28, 771–776.

Ren, G.R., Hauser, F., Rewitz, K.F., Kondo, S., Engelbrecht, A.F., Didriksen, A.K., Schjott, S.R., Sembach, F.E., Li, S., Sogaard, K.C., et al. (2015). CCHamide-2 Is an Orexigenic Brain-Gut Peptide in Drosophila. PLoS One 10, e0133017.

Robinson, M.D., McCarthy, D.J., and Smyth, G.K. (2010). edgeR: a Bioconductor package for differential expression analysis of digital gene expression data. Bioinformatics 26, 139–140.

Ruby, J.G., Stark, A., Johnston, W.K., Kellis, M., Bartel, D.P., and Lai, E.C. (2007). Evolution, biogenesis, expression, and target predictions of a substantially expanded set of Drosophila microRNAs. Genome research 17, 1850–1864.

Sano, H., Nakamura, A., Texada, M.J., Truman, J.W., Ishimoto, H., Kamikouchi, A., Nibu, Y., Kume, K., Ida, T., and Kojima, M. (2015). The Nutrient-Responsive Hormone CCHamide-2 Controls Growth by Regulating Insulin-like Peptides in the Brain of Drosophila melanogaster. PLoS Genet 11, e1005209.

Saper, C.B., and Fuller, P.M. (2017). Wake-sleep circuitry: an overview. Curr Opin Neurobiol 44, 186–192.

Shafer, O.T., and Keene, A.C. (2021). The Regulation of Drosophila Sleep. Curr Biol 31, R38–R49.

Shaw, P.J., Cirelli, C., Greenspan, R.J., and Tononi, G. (2000). Correlates of sleep and waking in Drosophila melanogaster. Science 287, 1834–1837.

Shcherbata, H.R., Ward, E.J., Fischer, K.A., Yu, J.Y., Reynolds, S.H., Chen, C.H., Xu, P., Hay, B.A., and Ruohola-Baker, H. (2007). Stage-specific differences in the requirements for germline stem cell maintenance in the Drosophila ovary. Cell Stem Cell 1, 698–709.

Sitaraman, D., Aso, Y., Jin, X., Chen, N., Felix, M., Rubin, G.M., and Nitabach, M.N. (2015a). Propagation of Homeostatic Sleep Signals by Segregated Synaptic Microcircuits of the Drosophila Mushroom Body. Curr Biol 25, 2915–2927.

Sitaraman, D., Aso, Y., Rubin, G.M., and Nitabach, M.N. (2015b). Control of Sleep by Dopaminergic Inputs to the Drosophila Mushroom Body. Front Neural Circuits 9, 73.

Song, Y., Ori-McKenney, K.M., Zheng, Y., Han, C., Jan, L.Y., and Jan, Y.N. (2012). Regeneration of Drosophila sensory neuron axons and dendrites is regulated by the Akt pathway involving Pten and microRNA bantam. Genes Dev 26, 1612–1625.

Sun, J., Xu, A.Q., Giraud, J., Poppinga, H., Riemensperger, T., Fiala, A., and Birman, S. (2018). Neural Control of Startle-Induced Locomotion by the Mushroom Bodies and Associated Neurons in Drosophila. Front Syst Neurosci 12, 6.

Ueno, K., Naganos, S., Hirano, Y., Horiuchi, J., and Saitoe, M. (2013). Long-term enhancement of synaptic transmission between antennal lobe and mushroom body in cultured Drosophila brain. J Physiol 591, 287–302.

Ueno, K., Suzuki, E., Naganos, S., Ofusa, K., Horiuchi, J., and Saitoe, M. (2017). Coincident postsynaptic activity gates presynaptic dopamine release to induce plasticity in Drosophila mushroom bodies. Elife 6.

Wang, Y., Mamiya, A., Chiang, A.S., and Zhong, Y. (2008). Imaging of an early memory trace in the Drosophila mushroom body. J Neurosci 28, 4368–4376.

Webb, J.M., and Fu, Y.H. (2020). Recent advances in sleep genetics. Curr Opin Neurobiol 69, 19–24.

Weng, R., and Cohen, S.M. (2015). Control of Drosophila Type I and Type II central brain neuroblast proliferation by bantam microRNA. Development 142, 3713–3720.

Xia, X., Fu, X., Du, J., Wu, B., Zhao, X., Zhu, J., and Zhao, Z. (2020). Regulation of circadian rhythm and sleep by miR-375-timeless interaction in Drosophila. FASEB J 34, 16536–16551.

Xie, X., Tabuchi, M., Corver, A., Duan, G., Wu, M.N., and Kolodkin, A.L. (2019). Semaphorin 2b Regulates Sleep-Circuit Formation in the Drosophila Central Brain. Neuron 104, 322–337 e314.

Yamazaki, D., Hiroi, M., Abe, T., Shimizu, K., Minami-Ohtsubo, M., Maeyama, Y., Horiuchi, J., and Tabata, T. (2018). Two Parallel Pathways Assign Opposing Odor Valences during Drosophila Memory Formation. Cell Rep 22, 2346–2358.

Yordanova, J., Kolev, V., Verleger, R., Bataghva, Z., Born, J., and Wagner, U. (2008). Shifting from implicit to explicit knowledge: different roles of early- and late-night sleep. Learn Mem 15, 508–515.

Zhang, R., Zhao, X., Du, J., Wei, L., and Zhao, Z. (2021). Regulatory mechanism of daily sleep by miR-276a. FASEB J 35, e21222.

